# Dynamic Functional Hyperconnectivity after Psilocybin Intake is Primarily Associated with Oceanic Boundlessness

**DOI:** 10.1101/2023.09.18.558309

**Authors:** Sepehr Mortaheb, Larry D. Fort, Natasha L. Mason, Pablo Mallaroni, Johannes G. Ramaekers, Athena Demertzi

## Abstract

To provide insights into neurophenomenological richness after psilocybin intake, we investigated the link between dynamical brain patterns and the ensuing phenomenological pattern after psilocybin intake. Healthy participants received either psilocybin (n=22) or placebo (n=27) while in ultra-high field 7T MRI scanning. Changes in the phenomenological patterns were quantified using the 5-Dimensional Altered States of Consciousness (5D-ASC) Rating Scale, revealing alterations across all dimensions under psilocybin. Changes in the neurobiological patterns displayed that psilocybin induced widespread increases in averaged functional connectivity. Time-varying connectivity analysis unveiled a recurrent hyperconnected pattern characterized by low BOLD signal amplitude, suggesting heightened cortical arousal. In terms of neurophenomenology, canonical correlation analysis primarily linked the transition probabilities of the hyperconnected pattern with feelings of oceanic boundlessness (OBN), and secondly with visionary restructuralization. We suggest that the brain’s tendency to enter a hyperconnected-hyperarousal pattern under psilocybin represents the potential to entertain variant mental associations. For the first time, these findings link brain dynamics with phenomenological alterations, providing new insights into the neurophenomenology and neurophysiology of the psychedelic state.

## Introduction

Hallucinogens are psychoactive drugs that, historically, have been used to alter conscious experience^1,2^. These drugs are divided into the classes of serotonergic psychedelics (e.g., psilocybin), antiglutamatergic dissociatives (e.g., ketamine), anticholinergic deliriants (e.g., scopolamine) and kappa-opioid agonists (e.g., salvinorin A)^3^. Research on classical hallucinogens has focused largely on serotonergic psychedelics, such as lysergic acid diethylamide (LSD), ayahuasca, psilocybin, N-dimethyltryptamine (DMT), and mescaline^4^. Among them, psilocybin has been one of the most studied psychedelics, possibly due to its potential contribution in treating different disorders^5^, such as obsessive-compulsive disorder^6^, death-related anxiety^7^, depression^8–11^, treatment-resistant depression^12–14^, major depressive disorder^15^, terminal cancer-associated anxiety^11,16^, demoralization^17^, smoking ^18^, and alcohol and tobacco addiction^19–21^.

The acute phase of psilocybin administration leads to the psychedelic state, which is a specific altered state of consciousness associated with consuming psilocybin and LSD^22^. By “state” we here refer to the combination of neurobiological and phenomenological patterns that are associated with the psychedelic experience.^24^ In terms of the general phenomenological pattern, the psychedelic state has been associated with the experience of ego dissolution (i.e., the reduction in self-referential awareness, ultimately disrupting self-world boundaries with increasing feelings of unity with others and own surroundings)^25,26^, unconstrained and hyper-associative cognition^27,28^, profound alterations in the perception of time, space and selfhood^29,30^, perceptual alterations, synesthesia, amplification of emotional state^31^, and emotional volatility^32^. Long-term and enduring effects have also been reported on personality and mood, such as increases in openness and extraversion, decreases in neuroticism, and increases in mindful awareness^33–35^. In terms of the neurobiological pattern, the psilocybin administration acutely resulted in increased global connectivity with reduced modularity^36–38^. Region-wise, there were reports of decreased activity in the thalamus, posterior cingulate cortex, medial prefrontal cortex^10^, and altered connectivity of the claustrum^39^. Network-wise, decreased connectivity was reported within the default mode network (DMN)^1,10^, visual network^1^, and executive control network (ECN)^40^, as well as reduced segregation of the dorsal attentional network and ECN^41^. These neural counterparts indicate that the subjective effects of psilocybin are linked to alterations in the activity and connectivity of important brain regions involved in information integration and routing when averaged signal analysis is concerned. Dynamic analyses of connectivity patterns after psilocybin administration have shown that the brain tended to recurrently configure into transient functional patterns with low stability^42^. In addition, under psilocybin, there were higher probabilities for the brain to configure into a connectivity pattern characterized by a global cortex-wide positive phase coherence^43^. In terms of state transition dynamics, a recent study calculated the minimum network control energy required to transition between states (or maintain the same state) and found that the network control energy landscape was flattened under LSD and psilocybin, meaning that there were more frequent state transitions and increased entropy of brain pattern dynamics^44^. Taken together, averaged and dynamic connectivity analyses suggest that psilocybin alters brain function, such that the overall neurobiological pattern becomes functionally more connected, more fluid, and less modular.

Of interest is the link between the psychedelic state’s neurobiological and phenomenological patterns as they are often analyzed separately. A recent investigation tried to correlate the occurrence rates of prominent connectivity patterns (i.e., frontoparietal subsystem and a globally coherent pattern) with subjective drug intensity (SDI) measured on a 10-point Likert scale^43,45^. As much as this approach has provided insights into the ensuing psychedelic state, it can be argued that, due to its simplicity, the SDI cannot capture the complexity of the psychedelic state’s phenomenological pattern. Here, we adopt a neurophenomenological approach after psilocybin intake to quantify its effects on cerebral functional dynamics and link these dynamic spatiotemporal fingerprints with reported experiential alterations measured with psychometric instruments.

## Results

Data was collected from 49 healthy participants who had previous experience with a psychedelic drug but not within the past three months^1^. Participants were allocated to two groups after being randomized to receive a single dose of psilocybin (0.17 mg/kg, n=22 (12 male, age=23±2.9 y) or placebo (bittering agent; n=27 (15 male, age=23.1±3.8 y). Six minutes of resting state with eyes open were acquired on a fMRI ultra-high field 7T scanner during the peak subjective drug effect (102 min post-treatment). The 5-Dimensional Altered States of Consciousness (5D-ASC) Rating Scale ^26,46^ was retrospectively evaluated at 360 min after drug administration.

### Psilocybin administration leads to a distinct phenomenological pattern

The psychedelic state’s phenomenological pattern was assessed using the 5D-ASC and its 11-ASC factors (see Methods) and were first assessed for normality assumptions. Shapiro-Wilk tests set at α=0.05 showed that all variables violated the assumption of normality (Table S1). As a result, Mann-Whitney U tests were used to compare the phenomenological outcomes in the two groups. Analyses revealed significant differences in all dimensions and factors with large effect sizes, such that the psilocybin group had more substantial effects than the placebo (Figure 1A and 1B, Table 1).

**Figure 1.**
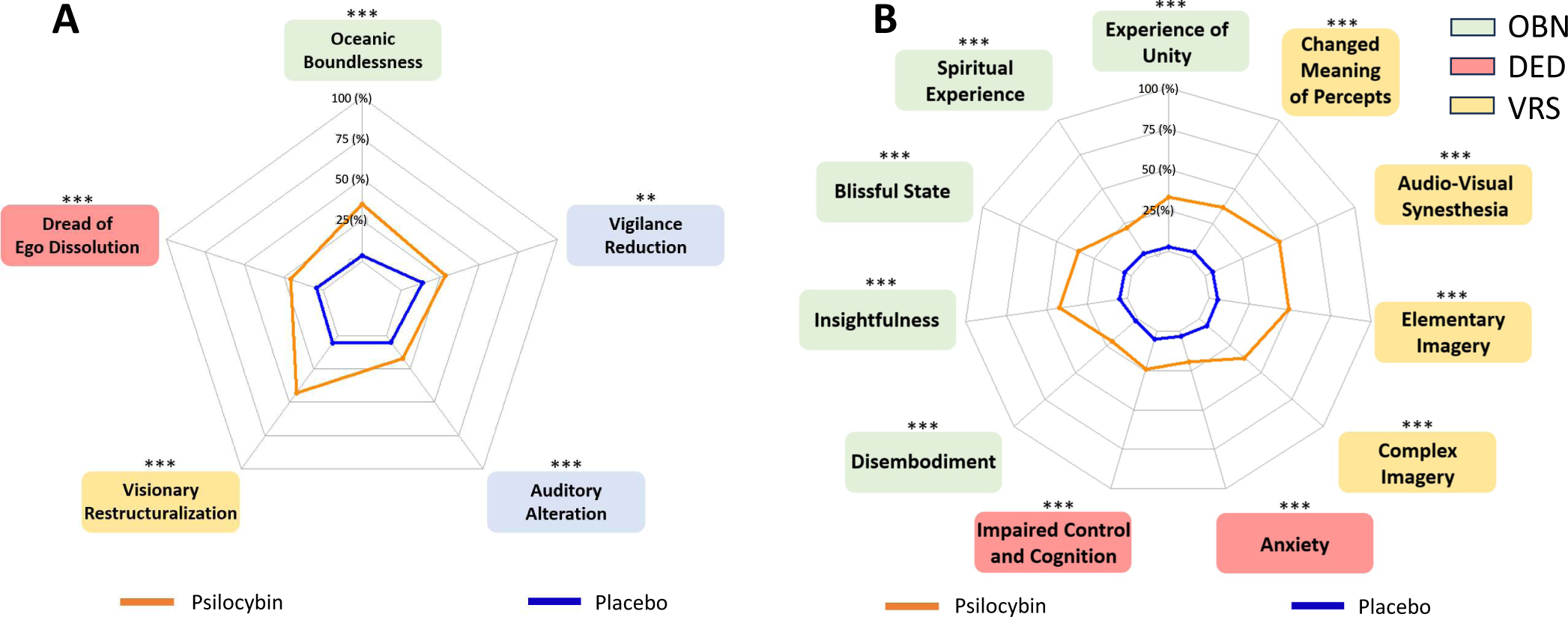
Substantial alterations in subjective experience were reported after psilocybin administration compared to placebo. A) The assessment of five dimensions of altered states of consciousness questionnaire (5D-ASC) showed that the administration of psilocybin significantly altered subjective experience in all dimensions. **B)** The same effect can also be observed considering the 11 factors of altered states of consciousness (11-ASC). Notes, OBN: Oceanic Boundlessness, DED: Dread of Ego Dissolution, VRS: Visionary Restructuralization. Radar plots illustrate group means for each dimension/factor.

**Table 1.** There were substantial changes in subjective experience after psilocybin administration compared to placebo. Mann-Whitney U test results of comparison between 11-ASC factors of the *Psilocybin* and *Placebo* groups. rg = Glass rank biserial coefficient effect size.

|  |  | Psilocybin<br>(n=21) |  | Placebo<br>(n=26) |  | Stats<br>(MW-U test) |  |  |
| --- | --- | --- | --- | --- | --- | --- | --- | --- |
|  |  | mean | SD | mean | SD | U | rg | p |
| 5D-ASC | Oceanic Boundlessness | 35.09 | 23.58 | 3.78 | 5.56 | 528.5 | .936 | < 0.001 |
|  | Dread of Ego Dissolution | 20.54 | 17.12 | 3.94 | 8.05 | 494 | .81 | < 0.001 |
|  | Visionary Restructuralization | 42.97 | 19.20 | 5.33 | 9.23 | 528.5 | .936 | < 0.01 |
|  | Auditory Alterations | 16.99 | 17.18 | 4.69 | 12.97 | 458.5 | .679 | < 0.001 |
|  | Vigilance Reduction | 28.18 | 20.43 | 13.99 | 16.33 | 400.5 | .467 | < 0.001 |
| 11-ASC | Insightfulness | 42.44 | 29.77 | 4.99 | 7.39 | 508 | 0.861 | <0.001 |
|  | Spiritual Experience | 21.73 | 22.04 | 3.33 | 6.91 | 435 | 0.593 | <0.001 |
|  | Experience of Unity | 33.27 | 30.57 | 2.85 | 4.70 | 483.5 | 0.771 | <0.001 |
|  | Blissful State | 35.65 | 27.02 | 4.36 | 7.04 | 489 | 0.791 | <0.001 |
|  | Disembodiment | 20.65 | 25.71 | 1.74 | 1.82 | 434 | 0.59 | <0.001 |
|  | Anxiety | 19.55 | 22.57 | 3.01 | 4.30 | 430 | 0.575 | <0.001 |
|  | Impaired Control and Cognition | 24.29 | 18.35 | 4.77 | 12.51 | 497 | 0.821 | <0.001 |
|  | Changed Meaning of Percept | 36.75 | 29.65 | 4.24 | 8.92 | 495.5 | 0.815 | <0.001 |
|  | Audio-Visual Synesthesia | 49.54 | 22.82 | 4.94 | 12.99 | 525 | 0.923 | <0.001 |
|  | Complex Imagery | 36.43 | 25.49 | 6.22 | 12.03 | 528.5 | 0.839 | <0.001 |
|  | Elementary Imagery | 49.25 | 21.95 | 5.68 | 9.24 | 502 | 0.943 | <0.001 |

### Functional connectivity and BOLD signal amplitude change after psilocybin administration

To investigate the neural effects of psilocybin, we first estimated the average functional connectome of each subject over the acquisition time. We used the Schaefer atlas to parcellate the brain into 100 distinct regions of interest (ROI) and computed Pearson correlations between pairs of ROIs. We found that after administration of psilocybin, whole-brain averaged connectivity increased (independent t-test: t=3.087, p=0.004; Figure 2A). This overall increase was further observed as a cortex-wide increase in the connectivity matrix values (independent t-test on the between-region connectivity values with FDR correction; Figure 2B). These alterations in the connectivity values were also accompanied by changes in the BOLD signal amplitude. By calculating the Euclidean norm of the BOLD time series related to each ROI, we found that regional BOLD signal amplitude decreased after psilocybin administration in both posterior and anterior regions compared to the placebo group (independent t-test, FDR-corrected; Figure 2C, and Figure S1). While somatomotor, limbic network and temporal regions of the DMN did not show significant changes in their signal norm, the highest decrease in BOLD signal amplitude was related to the posterior cingulate cortex and parietal regions of the ECN.

**Figure 2.**
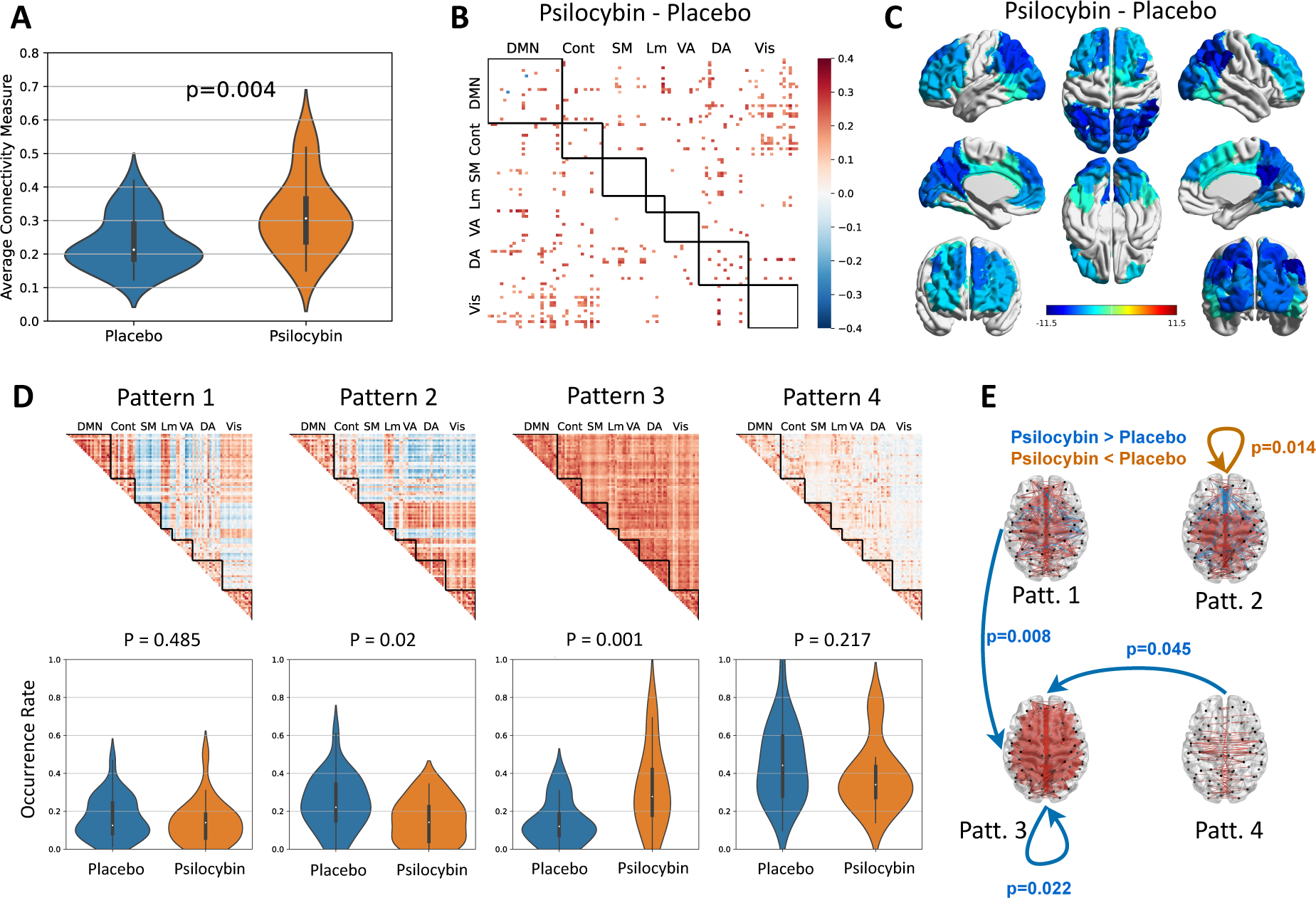
After psilocybin administration, there was an overall cerebral tendency to show more re-occurrence of a functional hyper-connectivity pattern. A) Averaged functional connectivity expressed as Fisher-transformed correlation values increased significantly after psilocybin administration compared to the placebo group. **B)** There were higher inter-regional connectivity values in the psilocybin group. The matrix represents the difference between the averaged connectivity matrix of the psilocybin group and that of the placebo group (contrast: psilocybin minus placebo). Only significant difference values are colored. **C)** The BOLD amplitude of posterior and anterior brain regions decreased after psilocybin administration, while the amplitude of somatomotor and limbic areas as well as the temporal regions of default more network remain unchanged. Colors are based on the difference values between the mean value of Euclidean norm of BOLD time series in the psilocybin group and the placebo group at each ROI; only significant difference values are colored. **D)** The functional connectome reconfigures in four connectivity patterns, ranging from complex inter-areal interactions (Pattern 1) to a low inter-areal connectivity profile (Pattern 4). After psilocybin administration, there was a significant increase in the occurrence rate of the global cortex-wide positive connectivity (Pattern 3). The connectivity matrices are colored based on the connectivity value: from dark blue to dark red corresponds to connectivity values from -1 to +1. Violin plots represent the distribution of patterns’ occurrence rates across participants. **E)** The transition probability from other patterns to Pattern 3 increased in the psilocybin group. Arrows indicate transitions between functional connectivity states. Blue corresponds to significantly higher transition probabilities (Wilcoxon Rank-Sum test) for the psilocybin group compared to the placebo, and orange corresponds to significantly higher transition probabilities for the placebo compared to the psilocybin group.

Next, we investigated the effect of psilocybin administration on dynamic changes of the whole-brain functional connectome by estimating phase-based coherence connectivity matrices at each time point of the ROI time series. After concatenating all connectivity matrices across participants, we applied K-means clustering to summarize them into four connectivity patterns (Figure 2D). Variant and distinct patterns of complex inter-areal interactions emerged: one of both correlations and anti-correlations (Pattern 1), one of anti-correlations of the DMN with other networks (Pattern 2), one of global cortex-wide positive connectivity (Pattern 3), and one of low inter-areal connectivity (Pattern 4). Pattern 3 occurred significantly more often in the psilocybin group when compared to the placebo group (independent t-test: t=3.731, p=0.001, α_*bonferroni*_ = 0.05/4 = 0.0125, Figure 2D). Furthermore, using Markov modeling and considering each of the four patterns as model states, we estimated the transition probabilities of each state to the others. The psilocybin group showed significantly higher transition probabilities towards Pattern 3 from Pattern 1 (Wilcoxon Rank-Sum test: z=2.744, p=0.006), Pattern 3 (z=2.291, p=0.022), and Pattern 4 (z=2.000, p=0.045; Figure 2E). In addition, the psilocybin group showed lower transition probabilities from Pattern 2 to itself compared to the placebo group (z=-2.452, p=0.014).

### The recurrent pattern of global hyperconnectivity is primarily associated with experiences of oceanic boundlessness and secondly with visionary restructuralization

To investigate the neurophenomenological counterpart of psilocybin intake, canonical correlation analysis (CCA) was performed between the 11-ASC factors and the pattern transition probabilities. CCA is a method of assessing the relationship between two multivariate datasets by maximizing their shared correlation while reducing the potential of type-I error^47^. For this purpose, we estimated the first canonical vector for both the behavioral and neural space that maximizes the shared correlation between two spaces (r=0.97, p<0.001). Considering the neural space, the transition probabilities from Pattern 1 to Pattern 3 showed the highest correlation with the first canonical vector of the neural space (r=0.86, p<0.001, Figure 3A). Transitions from Pattern 2 to Pattern 1 (r=0.47, p=0.015) and from Pattern 4 to Pattern 3 (r=0.40, p=0.048) showed lower significant correlations with the first canonical vector of the neural space. At the same time, in the phenomenological space, factors related to oceanic boundlessness (experience of unity: r=0.80, p<0.001, blissful state: r=0.74, p<0.001, insightfulness: r=0.68, p<0.001, spiritual experience: r=0.62, p<0.001, and disembodiment: r=0.35, p=0.036) and visionary restructuralization (elementary imagery: r=0.67, p<0.001, audio-video synesthesia: r=0.61, p<0.001, complex imagery: r=0.50, p=0.001, and changed meaning of percept: r=0.50, p=0.001) showed the highest correlations with the first canonical vector of the phenomenological space (Figure 3B). To validate the results, we performed another CCA analysis between the dimensions of the 5D-ASC and the between-state transition probabilities. At the neural space, the transition probability from Pattern 1 to Pattern 3 showed the highest correlation with the first canonical vector of the neural space (r=0.89, p<0.001), and at the phenomenological space, oceanic boundlessness showed the highest correlation with the first canonical vector of the phenomenological space (r=0.93, p<0.001; Table 2). These observations show that the experiences of oceanic boundlessness and visual restructuralization after psilocybin administration are primarily associated with the cerebral functional tendency to reconfigure into a global cortex-wide positive connectivity pattern.

**Figure 3.**
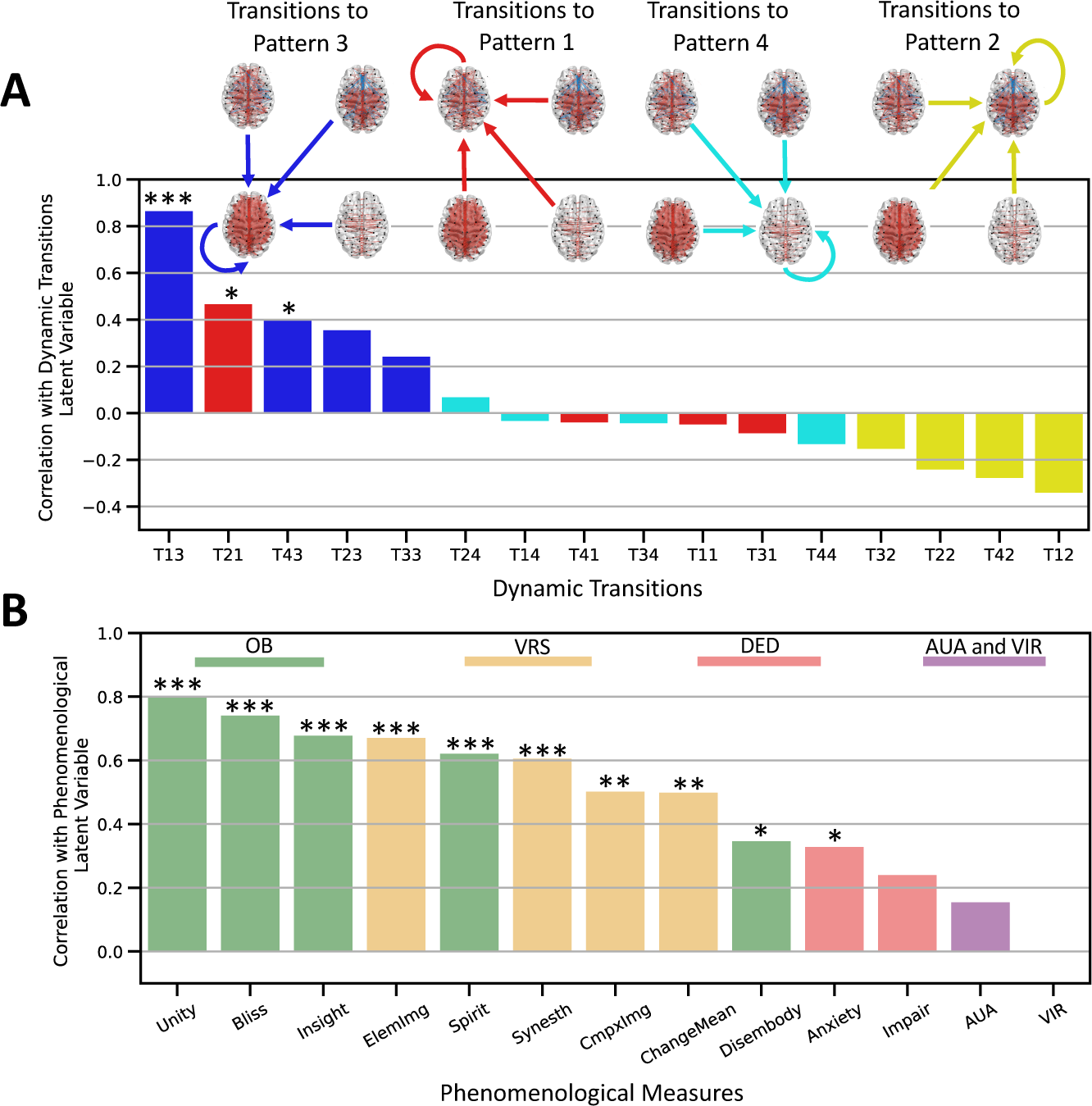
The neurophenomenological analysis indicated that transitions to the hyperconnected Pattern 3 were linked to the factors of oceanic boundlessness and visionary restructuralization. A) In the neural space, the canonical correlation analysis showed that the transition probabilities to the hyperconnected pattern had the highest correlation with the first canonical vector of the space. *Notes*: The y-axis represents pattern transitions, e.g. T13: transition from Pattern 1 to Patten 3. **B)** In the phenomenological space, factors related to the dimension of oceanic boundlessness and visionary restructuralization showed the highest correlation with the first canonical vector of the space. Bars represent correlation values of each factor to the first canonical vector of its associated space. Asterisks represent correlation significance (*p<0.05, **p<0.01, ***p<0.001). *Notes:* Unity: experience of unity, Bliss: blissful state, Insight: insightfulness, Spirit: spiritual experience, Disembody: disembodiment, ElemImg: elementary imagery, Synesth: audio-visual synesthesia, CmpxImg: complex imagery, ChangeMean: changed meaning of percept, Impair: Impaired control and cognition, AUA: auditory alterations, RIV: reduction of vigilance.

**Table 2.** The neurophenomenological analysis indicated that transitions from Pattern 1 to the hyperconnected Pattern 3 were linked to Oceanic Boundlessness. Canonical correlation analysis between between-state transition probabilities and five dimensions of the 5D-ASC. The results are sorted from the highest correlation to the lowest.

|  | Factor | Correlation Coefficient | p-value | p-value (FDR-Corrected) |
| --- | --- | --- | --- | --- |
| Neural Space | T13 | 0.89 | 1.54e-15 | 2.46e-14 *** |
|  | T23 | 0.44 | 3.66e-03 | 2.33e-02 * |
|  | T43 | 0.43 | 4.38e-03 | 2.33e-02 * |
|  | T33 | 0.25 | 1.12e-01 | 2.25e-01 |
|  | T21 | 0.23 | 1.52e-01 | 2.69e-01 |
|  | T14 | 0.12 | 4.36e-01 | 6.34e-01 |
|  | T32 | 0.09 | 5.65e-01 | 6.67e-01 |
|  | T24 | 0.09 | 5.89e-01 | 6.67e-01 |
|  | T31 | -0.01 | 9.41e-01 | 9.41e-01 |
|  | T44 | -0.08 | 6.26e-01 | 6.67e-01 |
|  | T11 | -0.09 | 5.63e-01 | 6.67e-01 |
|  | T41 | -0.17 | 2.92e-01 | 4.67e-01 |
|  | T34 | -0.27 | 7.86e-02 | 1.80e-01 |
|  | T22 | -0.33 | 3.13e-02 | 8.34e-02 |
|  | T12 | -0.34 | 2.93e-02 | 8.34e-02 |
|  | T42 | -0.34 | 2.85e-02 | 8.34e-02 |
| Phenomenological Space | OBN | 0.93 | 1.51e-19 | 7.55e-19 *** |
|  | VRS | 0.74 | 1.95e-08 | 4.89e-08 *** |
|  | DED | 0.46 | 1.94e-03 | 3.24e-03 ** |
|  | AUA | 0.35 | 2.46e-02 | 3.08e-02 * |
|  | VIR | 0.11 | 4.92e-01 | 4.92e-01 |
| Tij: probability of transition from Pattern i to Pattern j. OBN: oceanic boundlessness, VRS: visionary restructuralization, DED: dread of ego dissolution, AUA: auditory alteration, VIR: vigilance reduction. |  |  |  |  |

## Discussion

We investigated the effect of the serotonergic hallucinogen psilocybin on the brain’s functional connectome to link them with alternations of conscious experience to better comprehend how the resulting neural and experiential changes are interconnected. Overall, we found that psilocybin administration led to a tendency of the brain to recurrently configure in a globally hyperconnected pattern, which was linked to heightened reports of oceanic boundlessness (experience of unity, blissfulness, insightfulness, and spiritual experience), and visionary restructuralization (complex imagery, elementary imagery, audio-visual synesthesia, and changed meaning of percepts).

In terms of phenomenological changes, we report significant increases in all phenomenological dimensions and factors in the psilocybin group over the placebo. This includes increased reports of derealization, depersonalization, loss of self-control phenomena, elementary and complex visual pseudo-hallucinations, audio-visual synesthesia, experiences of unity, spiritual experience, bliss, insightfulness, and disembodiment^46^, as we previously reported^1^. These findings evidence that a moderate dose of psilocybin is enough to produce a significantly distinct phenomenological pattern when compared to a placebo. Previous research on psilocybin administration also found that dosages ranging from 45 to 315 μg/kg body weight produced measurable changes in oceanic boundlessness (OBN), visionary restructuralization (VRS), auditory alternations (AUA), vigilance reduction (VIR), and general altered states score (G-ASC) compared to placebo^48^. Additional research found that 260 μg/kg body weight of psilocybin produced significant changes in OBN, dread of ego dissolution (DED), and VRS when compared to placebo, ketanserin, and psilocybin with a pretreatment of ketanserin conditions^49^. Taken together, our data replicates the general, dose-dependent, phenomenological pattern associated with psilocybin consumption.

In terms of neural changes, we observed an overall increase in whole-brain functional connectivity in the psilocybin group, in line with previous reports^37,38^. Previous work showed that serotonergic psychedelics, including psilocybin, changed the brain’s functional organization into a new architecture, characterized by greater global integration, namely a higher amount of short-range and long-range functional connections^42,50^. This dynamic analysis evidenced that, under psilocybin, the brain had a higher probability of transitioning to a hyperconnected pattern compared to the placebo group. Similar configurations and dynamic transitions have been reported by other studies with psychedelic drugs, as well^43^. This hyperconnected pattern is characterized by maximal integration and minimal segregation^51^ and is interpreted as being functionally non-specific^43^. The significantly higher occurrence rate of this hyperconnected pattern in the psychedelic state can be explained by the “flattened landscape” theory^52^. Based on this dynamic systems approach, specific connectivity patterns, which act as attractors in typical conditions, become less dominant under psychedelics. Consequently, the brain expends less energy transitioning between these states^44^. This supports that the occurrence rate of functionally specific patterns in our analysis results in increased transition probabilities to enter the functionally non-specific hyperconnected pattern.

Moreover, we observed that this hyperconnectivity pattern is accompanied by cortex-wide decreases in the BOLD signal amplitude in the psilocybin group. We here decided to retain the global signal (GS) in the analysis pipeline, while acknowledging the significant debate around its removal as part of the denoising process^53^. On one hand, the GS has demonstrated a neuronal counterpart^54,55^., on the other, it was shown to reflect fMRI nuisance sources such as motion, scanner artifacts, respiration^56^, cardiac rate^57^, and vascular activity^58,59^. Our choice to keep the GS is justified by our recent finding that it can act as a complementary metric to extracted connectivity patterns^60^. Specifically, we found that when the hyperconnected pattern was accompanied by high GS amplitude during wakeful rest, it was more probable for participants to report phenomenological instances of mind blanking^60^ (i.e., the subjective evaluation of having no thoughts during unconstrained mentation^61^). The amplitude of the GS has been previously shown to act as an indirect measure of general arousal levels. For example, higher GS amplitude was related to lower levels of arousal, and lower GS amplitudes were linked to higher levels of arousal^62–65^. Indeed, it was demonstrated that the GS amplitude correlated positively with the relative amplitude of the delta band of EEG oscillations and negatively with the relative amplitude of the alpha band^64^. In the current study, we show that the hyperconnectivity pattern is accompanied by reduced GS amplitude, therefore adding to the explanation of the psychedelic state as mediated by high cortical arousal^66^.

In terms of neurophenomenology, we show that higher transition probabilities into the hyperconnected pattern are significantly associated with the factors of oceanic boundlessness (OBN) and visionary restructuralization (VRS). OBN is characterized by deeply felt positive mood, feelings of insight, and experiences of unity^26^. Previous studies associated positively experienced ego dissolution and oceanic boundlessness with reduced levels of hippocampal glutamate^1^ and higher feelings of insight with reduced levels of DMN within-network static functional connectivity^28^. Our whole-brain dynamic analysis complements this literature by showing that the brain’s tendency to be recurrently hyperconnected after psilocybin intake can also explain the experiences of unity. This is important as this factor is characterized by a disruption of the self- world boundary^1^ and the hyperconnected pattern is characterized by an atypical minimal segregation profile^51^. Other studies showed that both integration and segregation are necessary for a coherent sense of self^67^. Given these two main findings, we postulate that the reported feelings of unity and visual pseudo-hallucinatory experiences under psilocybin are connected to the brain’s inclination to exhibit highly integrated patterns, thereby evidencing the brain’s capacity to entertain diverse mental associations. This can be supported by the fact that different dimensions of creative thinking improve after psilocybin intake as a result of increased between-network functional connectivity of DMN and frontoparietal network,^28^ a connectivity profile also seen in the hyperconnected pattern.

Interestingly, three of the five factors of OBN (unity, bliss, insight) represented the highest canonical correlations in our analysis, while the visual restructuralization (VRS) factors followed. This was further supported by canonical correlations performed at the dimensional level, which showed OBN as having the highest association with transition probabilities to the hyperconnected pattern comparatively. This is a unique finding given that serotonergic psychedelics are sometimes referred to as hallucinogens, leaving one to assume that their primary effects are hallucinatory in nature^4^. Indeed, such an issue can be seen historically in the etymology of these drugs as they have been referred to as psychotomimetics, psycholytics, hallucinogens, psychedelics, or entheogens at different points in time^4,68^. Psilocybin-assisted therapy studies also show that the clinical outcome is driven primarily by OBN, supporting its primary role in the drug’s phenomenological effect^69^. OBN’s central importance is further supported by its association with decreased 5-HT2AR binding potential in cortical regions,^48,49^ its ability to predict medial prefrontal cortex-posterior cingulate cortex functional connectivity changes,^70^ and its positive correlation with disconnection of somatomotor network correlates^71^. Since OBN is defined as the positive valence associated with depersonalization and derealization through factors such as unity, that require dissolution of the self-world boundary, it can be seen primarily as a psychometric dimension that describes phenomenological modifications of the ego^26,46^. Our data, along with previous research, suggests that OBN is the primary driver of the psychedelic state’s general phenomenological pattern. As a consequence, future work might provide more refined terminologies to capture this driving effect of psilocybin, such as “egotropic” over “hallucinogenic”.

Our study is subject to various limitations. First, at the group level, the administration of psilocybin was not high enough to induce total ego dissolution for all participants. The administered dose was selected to be sufficient to induce a psychedelic state that participants could endure in an ultra-high field scanner environment. However, our phenomenological analysis showed that this dosage was adequate. Second, the lack of concurrent physiological recordings during fMRI scanning limits proper tracking of arousal levels. In this study, we used the GS amplitude as a proxy for the level of cortical arousal. At the same time, the absence of simultaneous electrophysiological recordings limits the direct interpretation of neuronal firing during hyperconnected patterns. Simultaneous physiological and electrophysiological recordings in future studies can help to better investigate the role of cortical and general arousal with neuronal firing patterns in producing mystical experiences during the psychedelic state. Finally, the data-driven nature of the analysis can limit the results of the current study to the recruited population. However, this is less concerning in the dynamic functional connectivity analysis as in our previous studies it was shown that similar recurrent connectivity patterns can be derived in different datasets and different brain parcellations, showing their replicability and universality^51,60^.

In conclusion, we found that pharmacological perturbations using psilocybin generate profound alterations both at the brain and at the experiential level. We showed that administration of psilocybin leads to an increase in the brain’s tendency to configure into a functionally non-specific hyperconnected organization, which phenomenologically associates with experiences of oceanic boundlessness and visual pseudo-hallucinations. The present findings illuminate the intricate interplay between brain dynamics and subjective experience under psilocybin, providing new insights into the neurophenomenology and neurophysiology of the psychedelic state.

## Methods

### Participants

Data were collected from 49 healthy participants with previous experience with a psychedelic drug, but not within the past 3 months of the experiment. Participants were randomized to receive a single dose of psilocybin (0.17 mg/kg, n=22; 12 male; age=23±2.9 y) or placebo (n=27; 15 male; age=23.1±3.8 y). This study was conducted according to the code of ethics on human experimentation established by the Declaration of Helsinki (1964) and amended in Fortaleza (Brazil, October 2013). This study is in accordance with the Medical Research Involving Human Subjects Act (WMO) and was approved by the Academic Hospital and the University’s Medical Ethics Committee (Maastricht University, Netherlands Trial Register: NTR6505). All participants were fully informed of all procedures, possible adverse reactions, legal rights, responsibilities, expected benefits, and their right to voluntary termination without consequences.

### Datasets

Six minutes of resting state fMRI were acquired from the participants with eyes open during peak subjective drug effect (102 minutes post-treatment). In addition, the 5-Dimensional Altered States of Consciousness (5D-ASC) Rating Scale was administered 360 minutes after drug intake, as retrospective phenomenological assessments of the drug experience.

### Phenomenological Assessment

The 5D-ASC Rating Scale is a 94-item self-report scale that assesses the participants’ alterations from normal waking consciousness^46^. In this questionnaire, participants are asked to make a vertical mark on the 10-cm line below each statement to rate the extent to which the following statements applied to their experience: “No, not more than usual” to “Yes, more than usual.” The 5D-ASC comprises five dimensions, including *oceanic boundlessness* (OBN), *dread of ego dissolution* (DED), *visionary restructuralization* (VRS), auditory *alterations* (AUA), and *vigilance reduction* (VIR). Further, OBN, DED, and VRS can be decomposed into 11 subscales via a previously published factor analysis^26^: OBN: *experience of unity*, *spiritual experience*, *blissful state*, *insightfulness*, *disembodiment*, DED: *impaired control and cognition*, *anxiety*, and VRS: *complex imagery*, *elementary imagery*, *audio-visual synesthesia*, and *changed meaning of percepts* comprising the 11-ASC scoring scheme.

### Imaging Setup

Images were acquired on a 7T Siemens Magnetom scanner (Siemens Medical, Erlangen, Germany) using 32 receiving channel head array Nova coil (NOVA Medical Inc., Wilmington MA). The T1w images were acquired using a magnetization-prepared 2 rapid acquisition gradient-echo (MP2RAGE) sequence collecting 190 sagittal slices following parameters: repetition time (TR) = 4500 ms, echo time (TE) = 2.39 ms, inversion times TI1 /TI2 = 900/2750 ms, flip angle1 = 5°, flip angle2 = 3°, voxel size = 0.9 mm isotropic, bandwidth = 250 Hz/pixel. In addition, 258 whole-brain EPI volumes were acquired at rest (TR = 1400 ms; TE = 21 ms; field of view=198 mm; flip angle = 60°; oblique acquisition orientation; interleaved slice acquisition; 72 slices; slice thickness = 1.5 mm; voxel size = 1.5 × 1.5 × 1.5 mm).

### Phenomenological Analysis

5D-ASC and its 11-ASC factors were assessed for normality assumptions using Shapiro-Wilk tests set at α=0.05. Due to violations in normality, non-parametric Mann-Whitney U tests were performed to compare the 5D-ASC and 11-ASC scores between the two groups. P-values were corrected using the Bonferroni method with a significance level of set at α=0.05. The effect size was calculated based on Glass rank biserial coefficient (rg).

### Neuroimaging Data Preprocessing

fMRI data were preprocessed using a locally developed pipeline based on SPM12^72^. After susceptibility distortion correction and realignment, functional data were registered to the high-resolution T1 image, then normalized to the standard MNI space, and finally smoothed using a Gaussian kernel with a full width at half maximum (FWHM) of 6mm. After segmentation of the structural T1 image into grey matter (GM), white matter (WM), and CSF masks, the bias-corrected structural image and all the extracted masks were normalized to the MNI space. Further, WM and CSF masks were eroded by one voxel to remove any overlapping between these tissues and the GM voxels. To denoise functional time series, we used a locally developed pipeline written in Python [nipype package^73^]. In this pipeline, a general linear model (GLM) was fitted to each voxel data separately, regressing out the effect of six movement parameters (translation in x, y, and z directions, and rotation in yaw, roll, and pitch directions) and their first derivative, constant and linear trends using zero-order and first-order Legendre polynomials, 5 principal components of signals in the WM and CSF masks, and outlier data points. Outlier detection was performed using the ART toolbox (http://web.mit.edu/swg/software.htm). Any volume with a movement value of greater than 3 mm, rotation value of greater than 0.05 radians, and z-normalized global signal intensity of greater than 3 was considered an outlier. After regressing out these nuisance regressors, the remaining signal was filtered in the range of [0.008, 0.09] Hz and was used for further analysis. The Schaefer atlas with 100 ROIs and a resolution of 2mm^74^ was used to extract the averaged BOLD signals inside each ROI.

### Averaged Functional Connectivity

Pearson correlations were calculated between the BOLD time series for every pair of ROIs, subsequently Fisher transformed, resulting in the generation of a 100×100 connectivity matrix for each individual participant. An Independent t-test was used to compare the 4950 possible between-region connectivity values between the two groups. FDR correction was performed to correct for multiple comparisons. Further, the average of the connectivity values over the whole brain was calculated for each participant and was considered as the overall connectivity value of the brain. An independent t-test was performed to compare the overall connectivity values between psilocybin and placebo groups.

### Time-varying Functional Connectivity

We used phase-based coherence analysis to extract between-region connectivity patterns at each time point of the scanning session^51,75^. For each participant i, after z-normalization of time series at each region r (i.e., xi,r[t]), the instantaneous phase of each time series was calculated via Hilbert transform as:

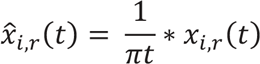

where * indicates a convolution operator. Using this transformation, we produced an analytical signal for each regional time series as:

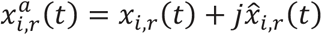

where 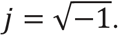 From this analytical signal, the instantaneous phase of each time series can be estimated as:

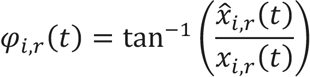

After wrapping each instantaneous phase signal of *φ*_*i,r*_ to the [−*π*, *π*] interval and naming the obtained signal as *θ*_*i,r*_(*t*), we calculated a connectivity measure for each pair of regions as the cosine of their phase difference. For example, the connectivity measure between regions r and s in subject i was defined as:

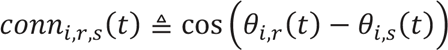

By this definition, completely synchronized time series lead to a connectivity value of 1, completely desynchronized time series produce a connectivity value of zero, and anticorrelated time series produce a connectivity measure of -1. Using this approach, we created a connectivity matrix of 100×100 at each time point t for each subject i that we called *C*_*i*_(*t*):

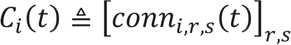

After collecting the connectivity matrices across all time points and participants, k-means clustering was applied, with 500 repetitions and 200 iterations at each repetition. With this technique, four robust and reproducible patterns were extracted as the centroids of the clusters, and each resting connectivity matrix was assigned to one of the extracted patterns.

We calculated the occurrence rate of each pattern defined as the proportion of connectivity matrices assigned to that pattern and was calculated for each subject separately. Independent two-tailed t-tests were used to compare the occurrence rate of each FC pattern between the psychedelic and placebo groups. Bonferroni correction was used to correct the p-values for multiple comparisons across the four connectivity patterns.

### Dynamic state transition modeling

To investigate the temporal evolution of the identified connectivity matrices, we defined the extracted patterns as the distinct states of a dynamical system transitioning between them over time using Markov modelling^51^. Using this approach, the data of a sample participant could be stated as a sequence of connectivity patterns over time (i.e., {*P*_*t*_ | *t*: 1, …, *T and P*_*t*_ ∈ {1, …, *M*}}, where M is the number of patterns and T is the number of signal time points). In this case, the probability of transitioning from Pattern *I* to Pattern *J* defined as *p*(*I* → *J*), considering *I*, *J* ∈ {1, . ., *M*}, can be calculated as the number of consecutive *I, J* pairs in the sequence, divided by the total number of transitions from pattern *I*:

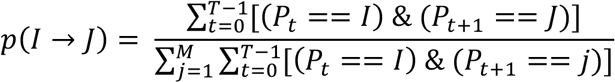

This transition probability was estimated for each possible between-state transition and each subject separately. With this approach, we could compare any significant difference in transition probabilities between two groups of subjects. To detect significantly different transition probabilities between the two groups, Wilcoxon rank-sum test was performed on each transition and p-values were FDR-corrected.

### Regional BOLD Amplitude Analysis

The Euclidean norm of the BOLD signal was calculated at each ROI as a measure of the power of the signal. Independent t-test was used to compare the regional GS power between two groups and p-values were FDR-corrected due to multiple comparisons.

### Neurophenomenological Analysis

A canonical correlation (CCA) analysis was conducted using the sixteen dynamic patterns transition probability variables as features of the neuronal space, and the 11-ASC and EDI variables as features of the phenomenological space, to evaluate the multivariate shared relationship between the two variable sets. CCA is a multivariate latent variable model that identifies associations between two different data modalities^47^. Considering matrix *X*^6×4^ contains M neuronal features of N subjects, and *Y*^6×8^ contains P phenomenological features of N subjects, the objective of the CCA is to find pairs of neuronal and phenomenological weights *W*^*M×1*^_*x*_ and *W*^*p×1*^_*x*_ such that the weighted sum of the neuronal and phenomenological variables maximizes the correlation between the resulting latent variables (canonical variates):

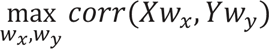

After finding latent variables *Xw*_*x*_ and *Yw*_*y*_ that have the maximal correlation, the features in each modality of data that have a stronger correlation with their respective latent variable are also significantly associated with one another.

## Data availability

The connectomes and the accompanying covariates used to differentiate individuals can be made available to qualified research institutions upon reasonable request to J.G.R. and a data use agreement executed with Maastricht University.

## Code availability

All codes used for analysis are freely available at https://gitlab.uliege.be/S.Mortaheb/psychedelics

## Author contributions

JR & AD contributed to the conception and design of the work; NM acquired the data; SM & LDF contributed with data analysis; all authors contributed to data interpretation; SM, LF, & AD drafted and revised the manuscript; all authors proofread the submitted work.

## Supporting information

Supplementary Indormation

## Acknowledgments

This article was supported by the Belgian Fund for Scientific Research (FRS-FNRS), the European Union’s Horizon 2020 Research and Innovation Marie Skłodowska-Curie RISE program NeuronsXnets (grant agreement 101007926), the European Cooperation in Science and Technology COST Action (CA18106), the Léon Fredericq Foundation, and the University and of University Hospital of Liège.

## Notes

### Competing Interest Statement

The authors have declared no competing interest.

### Summary of Updates

The discussion of the manuscript has been revised; Table 2 has been added to extend the CCA analysis results;

