## Supplementary Indormation for "Dynamic Functional Hyperconnectivity after Psilocybin Intake is Primarily Associated with Oceanic Boundlessness"

#### Supplementary Information

### Altered subjective experience after psilocybin intake associates with a dynamic pattern of hyperconnected functional connectivity


Sepehr Mortaheb[1,2], Larry D. Fort[1,2], Natasha L. Mason[3], Pablo Mallaroni[3], Johannes G. Ramaekers[3*±], Athena Demertzi[1,2,4*±]

[±]Equal contribution

[1]Physiology of Cognition, GIGA-CRC In Vivo Imaging, University of Liège, Belgium

[2]Fund for Scientific Research FNRS, Brussels, Belgium

[3]Department of Neuropsychology and Psychopharmacology, Faculty of Psychology and Neuroscience, Maastricht University, The Netherlands

[4]Psychology & Neuroscience of Cognition (PsyNCog), University of Liège, Belgium

[*]Corresponding authors:

**Table S1.** Shapiro-Wilk test results showed that the normality assumption is rejected for all the phenomenological variables. (OBE: oceanic boundlessness, VRS: visual restructuralization, DED: dread of ego dissolution, AUA: auditory alterations, VIR: vigilance reduction).

| | Variable | Statistic (W) | P-Value |
|---|---|---|---|
| **5D-ASC** | **OBE** | 0.762 | 1.58e-7 |
| | **VRS** | 0.826 | 4.48e-6 |
| | **DED** | 0.712 | 1.74e-8 |
| | **AUA** | 0.637 | 9.44e-10 |
| | **VIR** | 0.896 | 0.0004 |
| **11-ASC** | **Experience of Unity** | 0.659 | 2.06e-9 |
| | **Spiritual Experience** | 0.647 | 1.31e-9 |
| | **Blissful State** | 0.750 | 9.07e-8 |
| | **Insightfulness** | 0.741 | 6.03e-8 |
| | **Disembodiment** | 0.534 | 3.00e-11 |
| | **Complex Imagery** | 0.794 | 7.80e-7 |
| | **Elementary Imagery** | 0.814 | 2.23e-6 |
| | **Audio-Visual Synesthesia** | 0.743 | 6.58e-8 |
| | **Changed Meaning of Percepts** | 0.728 | 3.39e-8 |
| | **Impaired Control and Cognition** | 0.730 | 3.67e-8 |
| | **Anxiety** | 0.606 | 3.09e-10 |

**Figure S1**. Results of the independent t-test between regional Euclidean norm of the BOLD time series of *Psychedelic* and *Placebo* groups. Statistics (t-value) show the differences between the two groups (Placebo - Psilocybin) after FDR correction across the number of regions. The dashed line shows the significance threshold after FDR correction. T-values higher than the threshold are related to the significant difference.

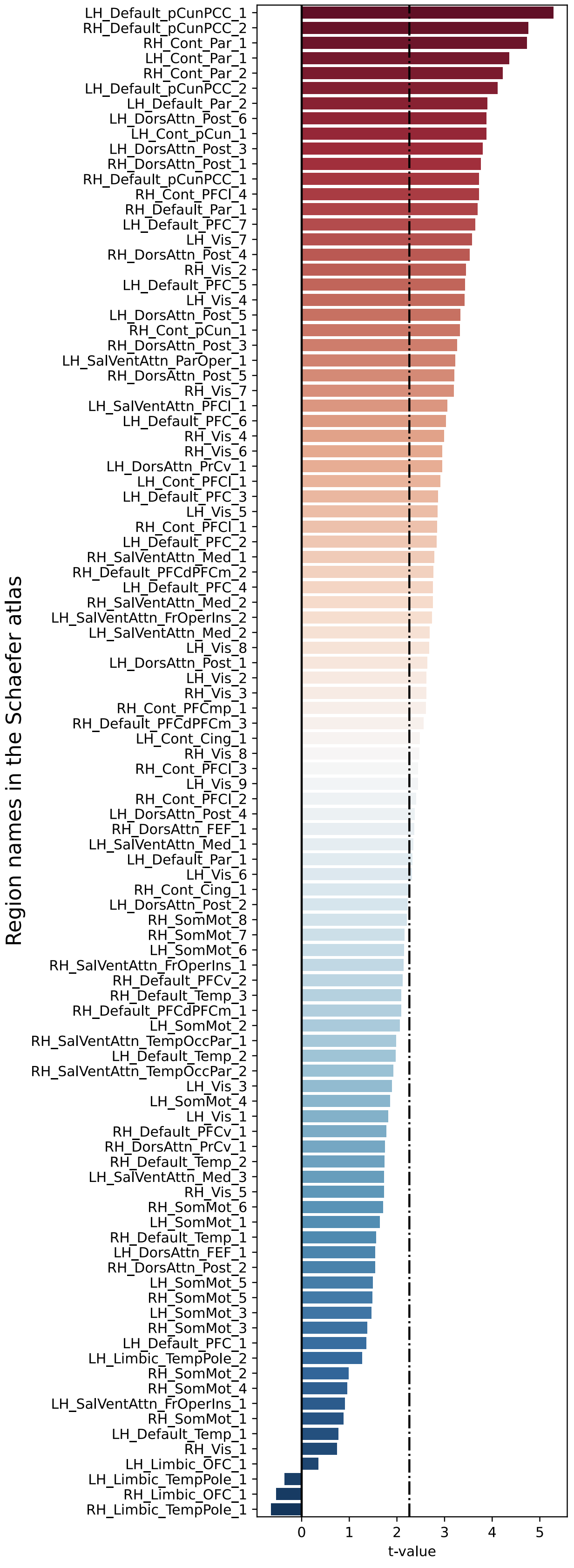
